# sstar: A Python package for detecting archaic introgression from population genetic data with *S*\*

**DOI:** 10.1101/2022.03.10.483765

**Authors:** Xin Huang, Patricia Kruisz, Martin Kuhlwilm

## Abstract

*S*\* is a widely used statistic for detecting archaic admixture from population genetic data. Previous studies used freezing-archer to apply *S*\*, which is only directly applicable to the specific case of Neanderthal and Denisovan introgression in Papuans. Here, we implemented sstar for a more general purpose. Compared with several tools, including SPrime, SkovHMM, and ArchaicSeeker2.0, for detecting introgressed fragments with simulations, our results suggest that sstar is robust to differences in demographic models, including ghost introgression and two-source introgression. We believe sstar will be a useful tool for detecting introgressed fragments in various scenarios and in non-human species.

---

Admixture between populations is a topic of great interest (Fontsere et al. 2019). To detect archaic admixture from population genetic data, a statistic named *S*\* was introduced to search for patterns of variation and linkage expected in the case of introgression (Plagnol and Wall 2006). This statistic has been applied in subsequent studies in modern humans (Huerta-Sanchez et al. 2014; Vernot and Akey 2014; Vernot et al. 2016; Xu et al. 2017; Jacobs et al. 2019) as well as other organisms (Cong et al. 2016; Kuhlwilm et al. 2019). Although the *S*\* statistic is a powerful approach for detecting introgressed fragments without source genomes (Browning et al. 2018), there is no user-friendly and versatile package available. A previous implementation of *S*\* is freezing-archer (Vernot and Akey 2014; Vernot et al. 2016), which was specifically designed with human demographic models and used for detecting introgressed fragments from Neanderthals and Denisovans into Papuans (Vernot et al. 2016). Users must carefully read and understand the source codes of freezing-archer before manually changing the parameters inside the code. To improve the efficiency, robustness and reproducibility when using *S*\* for detecting introgression, we implemented sstar.

The whole pipeline is illustrated in Figure 1A. We define the population without introgressed fragments as the reference population, the population which received introgressed fragments as the target population, and the population which donated introgressed fragments as the source population (Supplementary Figure S1). We assume genotype data are diploid, biallelic, phased and not missing in all individuals of the dataset. We also remove variants with derived alleles that are fixed in both the reference and target populations. Users can calculate *S*\* for sliding windows across genomes by defining the length of a window and step size. To assess significance of *S*\* scores, users can simulate data under a demographic model without introgression and build up a generalized linear model with different *S*\* scores, quantiles of *S*\*, numbers of mutations, and local recombination rates to predict the expected *S*\* score, as described previously (Vernot et al. 2016). If a genome from a potential source population is available, users can calculate the source match rate between an individual from the target population and an individual from the source population. If genomes from two different source populations are available, the origin of candidate introgressed fragments can be determined by comparing the source match rates with different source populations.

**Figure 1.**
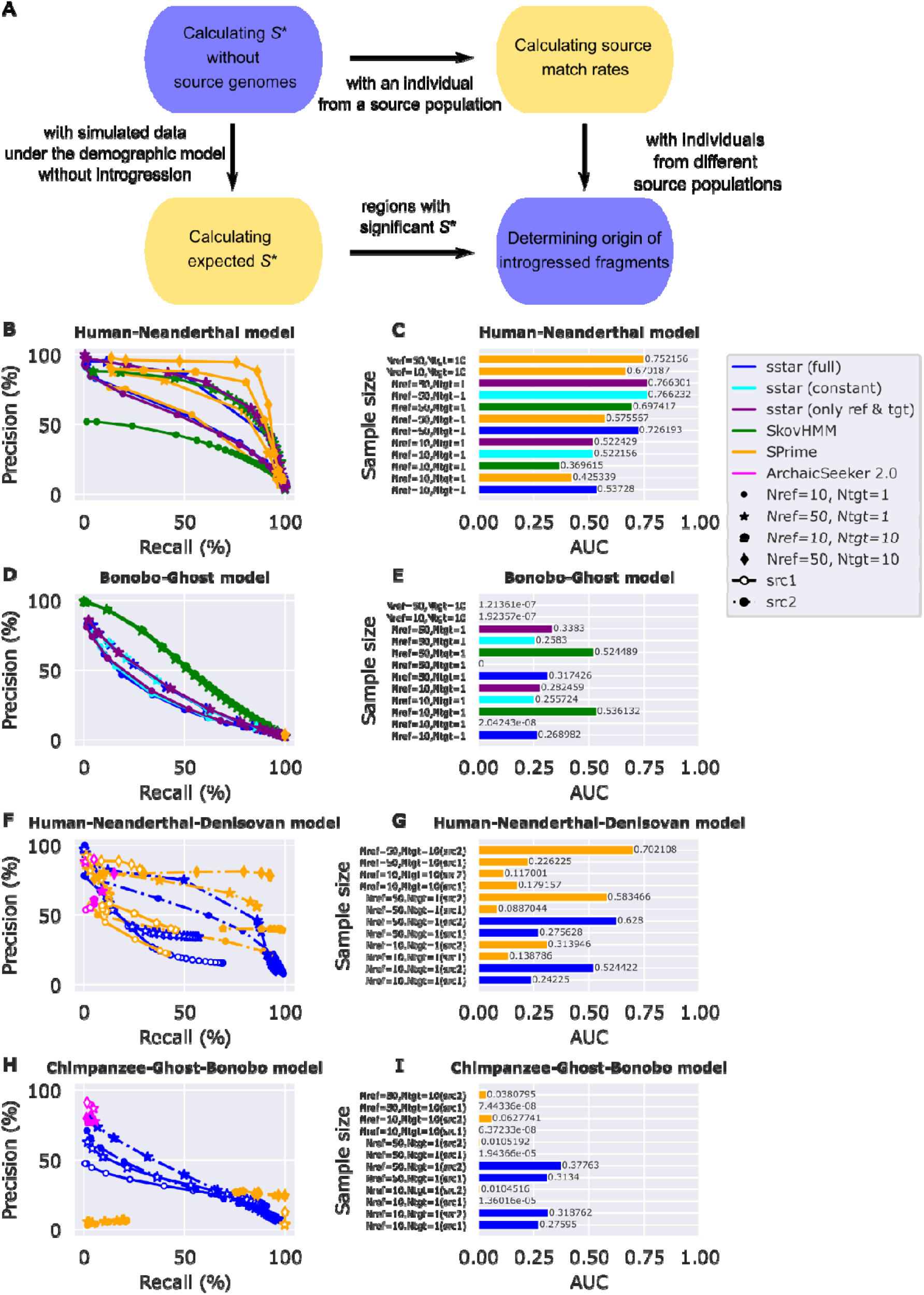
sstar workflow and its performance in different demographic models and sample sizes with SPrime, SkovHMM, and ArchaicSeeker2.0. Different points represent precision and recall estimated with different cut-offs (Supplementary Table S1–S6). Area under the curve (AUC) for each precision-recall curve was also estimated (Supplementary Table S7–S8). For ArchaicSeeker2.0, we only used the best results inferred from ArchaicSeeker2.0 and did not estimate AUC for ArchaicSeeker2.0, because ArchaicSeeker2.0 does not provide options to define candidate introgressed fragments with different cut-offs. For each demographic model, we simulated a sequence with a length of 200 Mb and replicated 100 times, then estimated the average performance of tools across different replicates. Nref is the diploid sample size of the reference population. Ntgt is the diploid sample size of the target population. For sstar and SkovHMM, we only used simulated data with 1 diploid individual from the target population, because these two tools analyse data individual by individual. sstar (full) are results inferred with generalized linear models (GLMs) using simulated data from full demographic models without introgression (Supplementary Figure S6–S9). sstar (constant) are results inferred with GLMs using simulated data from constant size models without introgression (Supplementary Figure S10 and S11). sstar (only ref & tgt) are results inferred with GLMs using simulated data from models with only the reference and target populations, these populations are also constant (Supplementary Figure S12 and S13). src1 represents the performance for identifying the introgressed fragments from the source population 1. src2 represents the performance for identifying the introgressed fragments from the source population 2. **(A)** The sstar workflow. *S*\* scores can be estimated without source genomes. Expected *S*\* scores can be obtained from simulated data without introgression by constructing GLMs and regions with significant *S*\* can be determined. If a source individual is available, source match rates can be estimated. If individuals from different source populations are available, the origin of the introgressed fragments can be determined by comparing source match rates estimated from different source individuals. **(B)** Precision-recall curves of sstar, SPrime, and SkovHMM for detecting introgression without source genomes under a Human-Neanderthal model (Gower et al. 2021, Supplementary Figure S2). **(C)** AUC scores for sstar, SPrime, and SkovHMM for detecting introgression without source genomes under a Human-Neanderthal model (Supplementary Figure S2). **(D)** Precision-recall curves of sstar, SPrime, and SkovHMM for detecting introgression without source genomes under a Bonobo-Ghost model (Kuhlwilm et al. 2019, Supplementary Figure S3). **(E)** AUC scores for sstar, SPrime and SkovHMM for detecting introgression without source genomes under a Bonobo-Ghost model (Supplementary Figure S3). **(F)** Precision-recall curves of sstar, SPrime, and ArchaicSeeker2.0 for detecting introgression with source genomes from two source populations under a Human-Neanderthal-Denisovan model from stdpopsim (Adrion et al. 2020, Supplementary Figure S4). The src1 population is the Neanderthal population. The src2 population is the Denisovan population. **(G)** AUC scores for sstar and SPrime for detecting introgression with source genomes from two source populations under a Human-Neanderthal-Denisovan model (Supplementary Figure S4). **(H)** Precision-recall curves of sstar, SPrime, and ArchaicSeeker2.0 for detecting introgression with source genomes from two source populations under a Chimpanzee-Ghost-Bonobo model (Supplementary Figure S5) modified from Kuhlwilm et al. (2019). The src1 population is the Ghost population, the src2 population is the Bonobo population, both introgressing into the Central Chimpanzee population. **(I)** AUC scores for sstar and SPrime for detecting introgression with source genomes from two source populations under a Chimpanzee-Ghost-Bonobo model (Supplementary Figure S5).

We evaluated the performance of sstar by simulation with msprime 1.0 (Kelleher et al. 2016; Baumdicker et al. 2022) in different demographies and sample sizes. Two models tested ghost introgression: a Human-Neanderthal model (Gower et al. 2021) and a Bonobo-Ghost introgression model (Kuhlwilm et al. 2019). Two models tested two-source introgression: a Human-Neanderthal-Denisovan model (Malaspinas et al., 2016; Jacobs et al., 2019) and a Chimpanzee-Ghost-Bonobo model. For ghost introgression, we compared sstar with SPrime (Browning et al. 2018), another tool using an *S*\*-like approach, and SkovHMM (Skov et al. 2018), a tool based on the hidden Markov model (HMM). In the Human-Neanderthal model, our results show that sstar performed the best among these approaches, when detecting introgression with 10 diploid reference individuals and 1 target individual (Figure 1B and 1C). In the Bonobo-Ghost model, SPrime performed poorly (Figure 1D and 1E), assigning the whole sequence as introgressed. In this model, sstar and SkovHMM still could detect introgressed fragments.

One key step in sstar is calculating the expected *S*\* scores with simulated data from demographic models without introgression, requiring full knowledge on the population history (Supplementary Figure S6–S9). Using approximate history (Supplementary Figure S10–S13), our results suggest that sstar performed similarly when compared with those using the full history (Figure 1B–1E). For two-source introgression, we compared sstar with SPrime, and ArchaicSeeker2.0 (Yuan et al. 2021; Zhang et al. 2022), another HMM-based tool. In the Human-Neanderthal-Denisovan model with large datasets, SPrime performed well, though both sstar and SPrime performed better when identifying Denisovan fragments than identifying Neanderthal fragments (Figure 1F and 1G). This may be due to the Denisovan introgression event in Papuans being more recent and its admixture proportion being larger than for the Neanderthal introgression. More ancient events like in the Chimpanzee-Ghost-Bonobo model cannot be well determined by SPrime, while sstar still retained power (Fig. 1H and 1I).

We conclude that sstar is robust for detecting introgressed fragments. Since no single tool could perform well for detecting introgressed fragments under different demographies and sample sizes, we believe sstar will be useful for exploring introgression in various scenarios, especially small samples, and non-humans.

## Data availability

Source codes for sstar can be found in https://github.com/admixVIE/sstar (last accessed on June 15, 2022) and the manual can be found in https://sstar.readthedocs.io/en/latest/ (last accessed on June 15, 2022). Codes for replicating the benchmark can be found in https://github.com/admixVIE/sstar-analysis (last accessed on June 15, 2022). Computational tools installed through conda can be found in https://github.com/admixVIE/sstar-analysis/blob/main/environment.yml (last accessed on June 15, 2022). Tools listed below cannot be installed through conda but can be found in the websites from their authors: ArchaicSeeker2.0 (https://github.com/Shuhua-Group/ArchaicSeeker2.0, last accessed on June 15, 2022), ms program (Hudson 2002; https://home.uchicago.edu/~rhudson1/source/mksamples.html, last accessed on June 15, 2022), SkovHMM (https://github.com/LauritsSkov/Introgression-detection, last accessed on June 15, 2022), SPrime (https://github.com/browning-lab/sprime, last accessed on June 15, 2022), and SPrime pipeline (Zhou and Browning 2021; https://github.com/YingZhou001/sprimepipeline, last accessed on June 15, 2022).

Demographic models in Demes YAML format (Gower et al. 2022) can be found in https://github.com/admixVIE/sstar-analysis/tree/main/config/simulation/models (last accessed on June 15, 2022).

## Acknowledgments

We thank Benjamin Vernot for discussions on freezing-archer, Andrew Kern and Peter Ralph for implementing Snakemake pipelines, Graham Gower for plotting demographic models with demesdraw and help from the PopSim Consortium. This project has been funded by the Vienna Science and Technology Fund (WWTF) and the City of Vienna through project VRG20-001.

## Author contribution

M.K. and X.H. designed the study. X.H. implemented sstar. M.K. tested sstar. X.H., P.K. and M.K. implemented the benchmark pipelines. X.H. and M.K. analysed the data and wrote the manuscript.

