## Supplementary Material for "sstar: A Python package for detecting archaic introgression from population genetic data with *S*\*"

**Brief Communication**


### Supplementary

### Materials and Methods

#### *Calculating S**

The *S** statistic was originally developed by Plagnol and Wall (2006). It captures introgressed fragments by measuring the level of linkage disequilibrium across genomes (Plagnol and Wall 2006). *S** can be estimated and used for detecting introgressed fragments without genomes from the source population. To efficiently calculate *S** across genomes, Vernot and Akey (2014) proposed a dynamic programming algorithm. Moreover, Vernot et al. (2016) extended *S** using a single diploid genome from the target population and a large panel of genomes from the reference population. In sstar, we calculate *S** as defined by Vernot et al. (2016) with the dynamic programming algorithm used by Vernot and Akey (2014).

If we have a genomic region with a length $L$ and $n$ variants, then we can define $V_k = \{1,2,\cdots,n\}$ as the set of variants located at this region in the *k*-th individual from the target population. The *S** of the *k*-th individual in this region is $S_k^* = S_k^*(H) = max_{H_k \subseteq V_k}\{S_k^*(H_k)\}$, where $H$ is a subset of $V_k$ maximizing $S_k^*$. We further define $d_{ij}$ and $g_{ij}$ as the physical and genotype distance between the $i$-th and $j$-th variants in a single individual, respectively. Because we assume biallelic variants and diploid individuals without missing data, $g_{ij} = \left|\sum_{l=1}^{2}(a_{il} - a_{jl})\right|, a_{il}, a_{jl} \in \{0,1\}$, where $a_{il}$ and $a_{jl}$ are the *l*-th alleles of the $i$-th and $j$-th variants, respectively. We then estimate $S_k^*$ with dynamic programming. Let $S_k^*(i)$ denote the $S$* for the $i$-th variant, and $H_k(i)$ denote the set of variants maximizing $S_k^*(i)$, then the $S$* for the $j$-th variant can be estimated as follow:

$$S_{k,i}^*(j) = max\left\{S_{k,i-1}^*(j), S_k^*(i,j), S_k^*(i) + S_k^*(i,j)\right\}, \qquad 1 \le i \le j$$

where $S_{k,i}^*(j)$ is the $S$* for the $j$-th variant after comparing the $j$-th variant with the $i$-th variant, $S_{k,0}^*(j) = -\infty$ and $S_k^*(j) = S_{k,j}^*(j)$. Here,

$$S_k^*(i,j) = \begin{cases} -\infty, & 0 \le d_{ij} < 10 \\ B + d_{ij}, & d_{ij} \ge 10 \text{ } and \text{ } g_{ij} = 0 \\ P, & d_{ij} \ge 10 \text{ } and \text{ } 0 < g_{ij} \le M \\ -\infty, & d_{ij} \ge 10 \text{ } and \text{ } g_{ij} > M \end{cases}$$

where $B$ is the bonus for matching two genotypes at the $i$-th and $j$-th variants, $P$ is the penalty for mismatching two genotypes at the $i$-th and $j$-th variants, and $M$ is the upper limit for mismatching two genotypes at the $i$-th and $j$-th variants. We do not estimate $S$* for two variants with a physical distance less than 10 base pairs (Plagnol and Wall 2006).

Also, for $1 \le i \le j$ , we have

$$H_{k,i}(j) = \begin{cases} H_k(i) \cup \{j\}, & S_{k,i}^*(j) = S_k^*(i) + S_k^*(i,j) \\ \{i,j\}, & S_{k,i}^*(j) = S_k^*(i,j) \\ H_{k,i-1}(j), & S_{k,i}^*(j) = S_{k,i-1}^*(j) \end{cases}$$

where $H_{k,i}(j)$ is the set of variants maximizing $S_{k,i}^*(j)$, $H_{k,0}(j) = \emptyset$ and $H_k(j) = H_{k,j}(j)$. If $S_{k,i}^*(j) = S_{k,i-1}^*(j) = S_k^*(i) + S_k^*(i,j)$, then $H_{k,i}(j) = H_{k,i-1}(j)$. A toy example for calculating $S$* with the above algorithm can be found in Vernot and Akey (2014).

***Calculating expected S* under the demographic model without introgression***

To determine whether an $S$* score is statistically significant, an expected $S$* score can be estimated using simulations, as described before (Vernot et al., 2016). In more detail, we simulate data using demographic models without introgression with a given length $L$, a given number of mutations $u$, and a given local recombination rate $r$. $L$ is equal to the length of the window used for detecting introgressed fragments, and $u$ and $r$ should represent the range of values observed in real data. We then retrieve simulated data multiple times (20,000 in our analysis) for each $u$ and $r$, then calculate $S$* for each replicate. Thus, we obtain the distribution of $S$* and different quantiles of $S$* for the given $u$ and $r$ values. A generalized linear model (GLM) is then built using the different numbers of mutations, different local recombination rates, and $S^*$ scores, and it can be determined for different quantiles of $S^*$. Using this generalized linear model, an $S^*$ score under the null model can be predicted by giving the observed number of mutations and local recombination rate in a region at a specific quantile. If the estimated $S^*$ score is larger than the $S^*$ score from the null model, we assign this region as a candidate introgressed fragment.

***Calculating source match rates with a single source individual***

If an individual from the source population is available, we also could estimate the similarity between a window from an individual the target population and the source individual. We define the source match rate of a window as

$$m = \frac{1}{2}\sum_{i=1}^{2}\frac{|M_i \cap N_{hom}| + 0.5|M_i \cap N_{het}|}{|M_i \cup N_{hom} \cup N_{het}|}$$

where $M_i$ is the set of variants found in the *i*-th haplotype of a diploid individual from the target population, $N_{hom}$ is the set of homozygous variants found in the source individual, and $N_{het}$ is the set of heterozygous variants found in the source individual (Vernot et al. 2016).

***Determining the origin of a candidate introgressed fragments***

If individuals from two different source populations are available, we could determine the origin of a candidate introgressed fragment by comparing the source match rates with different source populations. More specifically, if $m_1 - m_2 > d$, then we assign the candidate introgressed fragments to the source population 1. Here, $m_1$ is the source match rate estimated with the individual from the source population 1, and $m_2$ is the source match rate estimated with the individual from the source population 2. *d* is a user-defined threshold. If $m_1 - m_2 < d$, then we assign the candidate introgressed fragments to the source population 2. We used *d* = 0 in our analysis.

***Benchmarking***

We performed benchmarks for sstar, SPrime (version: 07Dec18.5e2), SkovHMM, and ArchaicSeeker2.0 on the Life Science Compute Cluster for the University of Vienna. We used msprime (version: 1.2.0) to simulate data from four demographic models (Supplementary Figure S2–S5). The parameters for these models can be found in Supplementary Table S9–S12. The first model was a Human-Neanderthal model (Supplementary Figure S2) based on the parameters from Gower et al. (2021). In this model, YRI was the reference population, and CEU was the target population. The second model was a Bonobo-Ghost model

(Supplementary Figure S3) based on the parameters from Kuhlwilm et al. (2019). In this model, the Western Chimpanzee population was the reference population, and the Bonobo population was the target population. The third model was a Human-Neanderthal-Denisovan model (Supplementary Figure S4) based on the parameters from the PapuansOutOfAfrica_10J19 model in stdpopsim. In this model, YRI was the reference population, and the Papuan population was the target population. Two source diploid individuals were sampled, one was from the Altai Neanderthal population, the other one was from the Altai Denisovan population. The fourth model was a Chimpanzee-Ghost-Bonobo model (Supplementary Figure S5) modified from the Bonobo-Ghost model. We removed the introgression event from the Ghost population to the Bonobo population and the migration events between the Central Chimpanzee population and the Bonobo population. Then we added two introgression events into the Central Chimpanzee population: one was from the Ghost population, the other one was from the Bonobo population. In this model, the Western Chimpanzee population was the reference population, and the Central Chimpanzee population was the target population. Two source diploid individuals were sampled, one was from the Ghost population, the other one was from the Bonobo population.

We used precision (ratio of amount of true introgressed fragments detected to amount of inferred introgressed fragments) and recall (ratio of true introgressed fragments detected to amount of true introgressed fragments) to measure the performance of each tool. When using sstar, we used sliding windows with a length of 50 kb and a step size of 10 kb across the genome in the Human-Neanderthal model and the Human-Neanderthal-Denisovan model, while we used sliding windows with a length of 40 kb and a step size of 10 kb across the genome in the Bonobo-Ghost model and the Chimpanzee-Ghost-Bonobo model. To construct GLMs for calculating expected $S^*$ scores without introgression, we used the ms program (Hudson 2002) to simulate data with fixed number of mutations (ranging from 25 to 700 with a step size of 5) in a window under different demographic models (Supplementary Figure S6–S13). We tested the performance of sstar with GLMs from simulated data using the full history without introgression (Supplementary Figure S6–S9) and approximate history without introgression (Supplementary Figure S10–S13). In the models of approximate history, we first assumed all the populations had the same population size 10,000 through time (Supplementary Figure S10–S11), then we further only used the reference and target populations with constant sizes to simulate data (Supplementary Figure S12–S13). In the ms simulation, we assume the mutation rate $1.29\times10^{-8}$ per base per generation and the recombination rate $1\times10^{-8}$ per base per generation for the Human-Neanderthal model, assume the mutation rate $1.4\times10^{-8}$ per base per generation and the recombination rate $1\times10^{-8}$ per base per generation for the Human-Neanderthal-Denisovan model, assume the mutation rate $1.2\times10^{-8}$ per base per generation and recombination rate $0.7\times10^{-8}$ per base per generation for the Bonobo-Ghost model and the Chimpanzee-Ghost-Bonobo model. When using SPrime, we set the mutation rate $1.29\times10^{-8}$ per base per generation and recombination rate $1\times10^{-8}$ per base per generation for the simulated data from the Human-Neanderthal model and set the mutation rate $1.4\times10^{-8}$ per base per generation and recombination rate $1\times10^{-8}$ per base per generation for the simulated data from the Human-Neanderthal-Denisovan model, while we set the mutation rate $1.2\times10^{-8}$ per base per generation and recombination rate $0.7\times10^{-8}$ per base per generation for the simulated data from the Bonobo-Ghost model and the Chimpanzee-Ghost-Bonobo model. We used the default values for other parameters in SPrime. In the Human-Neanderthal-Denisovan model and the Chimpanzee-Ghost-Bonobo model, we used the SPrime pipeline (Zhou and Browning 2021) to determine the origin of introgressed fragments by comparing the match rates estimated with different source populations. We assigned the candidate introgressed fragments to the source population with a large match rate. When using SkovHMM, we used non-overlapping windows with a length of 1000 bp across the genome. When using ArchaicSeeker2.0, we set the recombination rate $1\times10^{-8}$ per base per generation for the simulated data from the Human-Neanderthal model and the Human-Neanderthal-Denisovan model, while we set the recombination rate $0.7\times10^{-8}$ per base per generation for the simulated data from the Bonobo-Ghost model and the Chimpanzee-Ghost-Bonobo model. We used the default values for other parameters in ArchaicSeeker2.0.

### Supplementary Figures

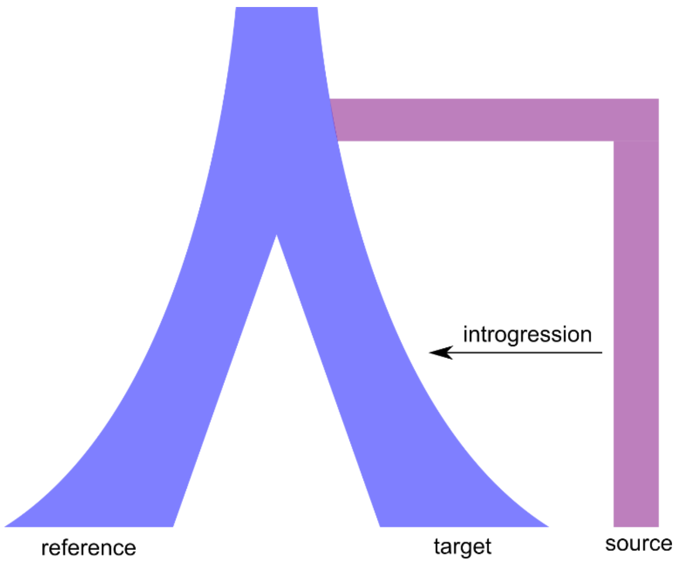


**Figure S1 Definition of populations in sstar.** The reference population is the population without introgressed fragments. The target population is the population that received introgressed fragments. The source population is the population that donated introgressed fragments to the target population.

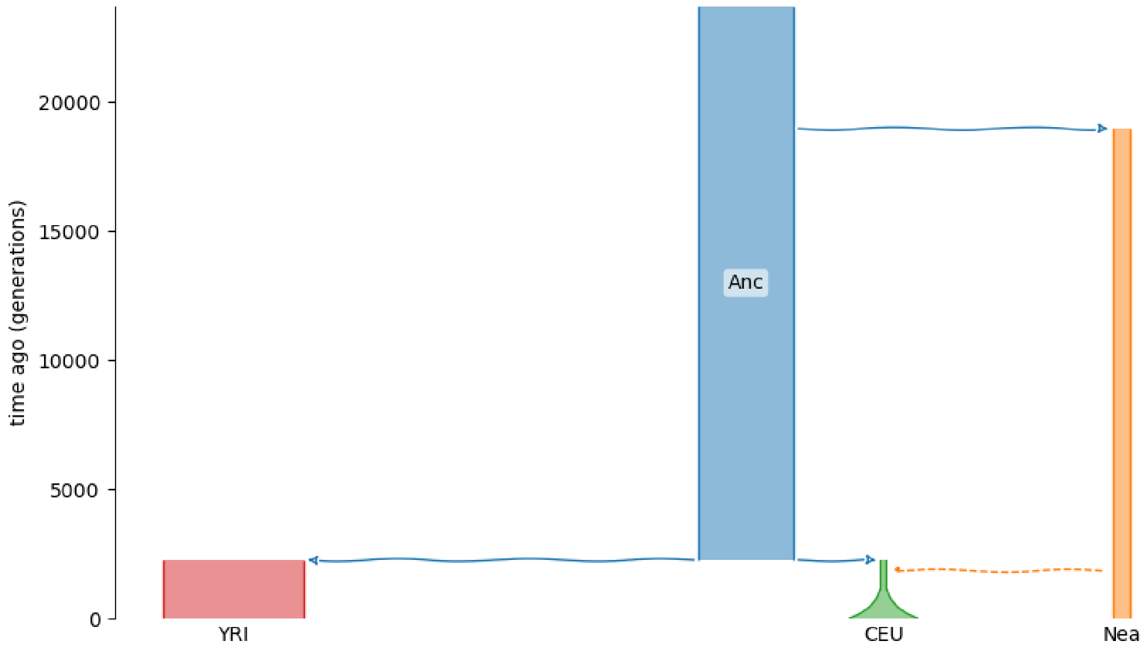


**Figure S2 Human-Neanderthal model for simulating data.** The demographic parameters can be found in Supplementary Table S9.

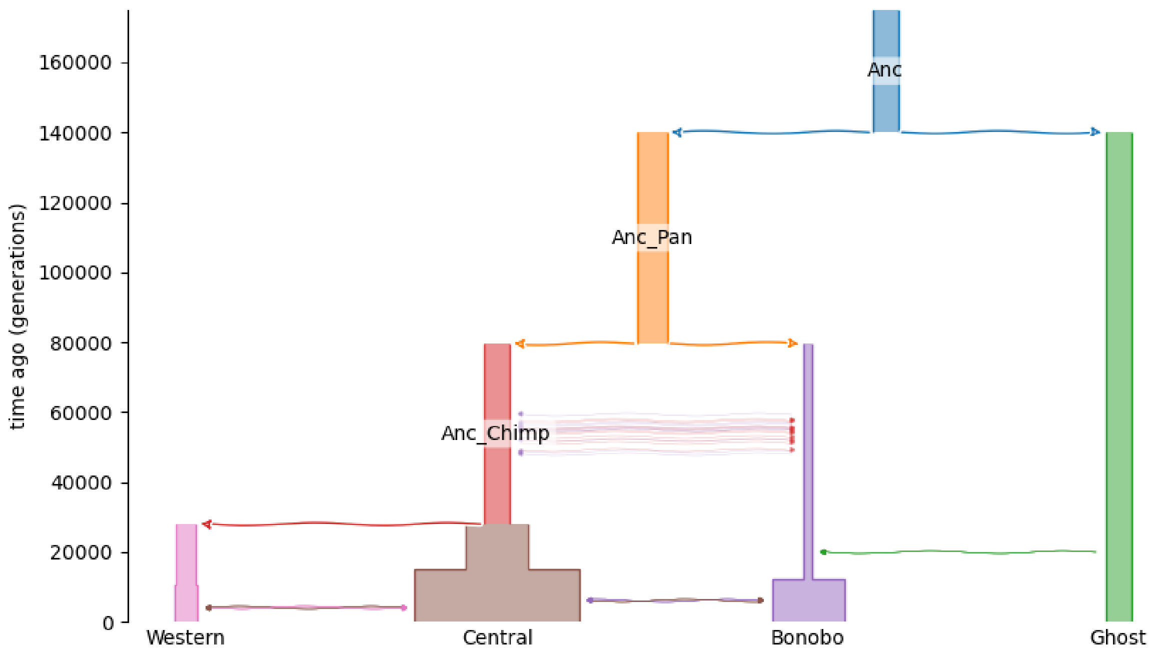


**Figure S3 Bonobo-Ghost model for simulating data.** The demographic parameters can be found in Supplementary Table S10.

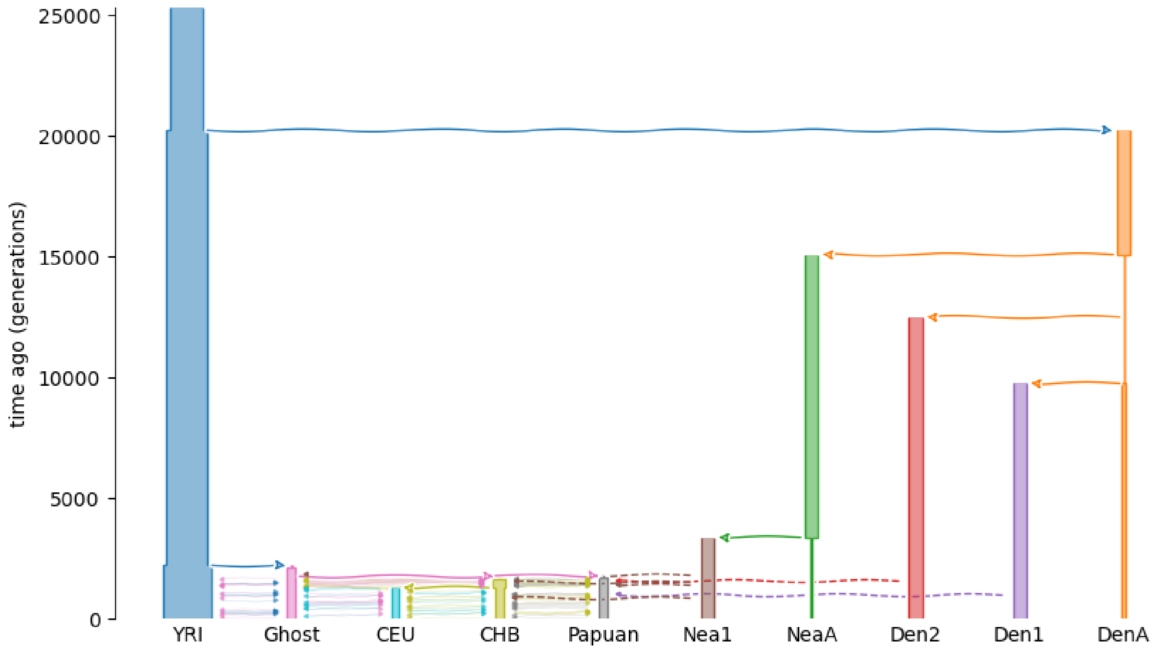


**Figure S4 Human-Neanderthal-Denisovan model for simulating data.** The demographic parameters can be found in Supplementary Table S11.

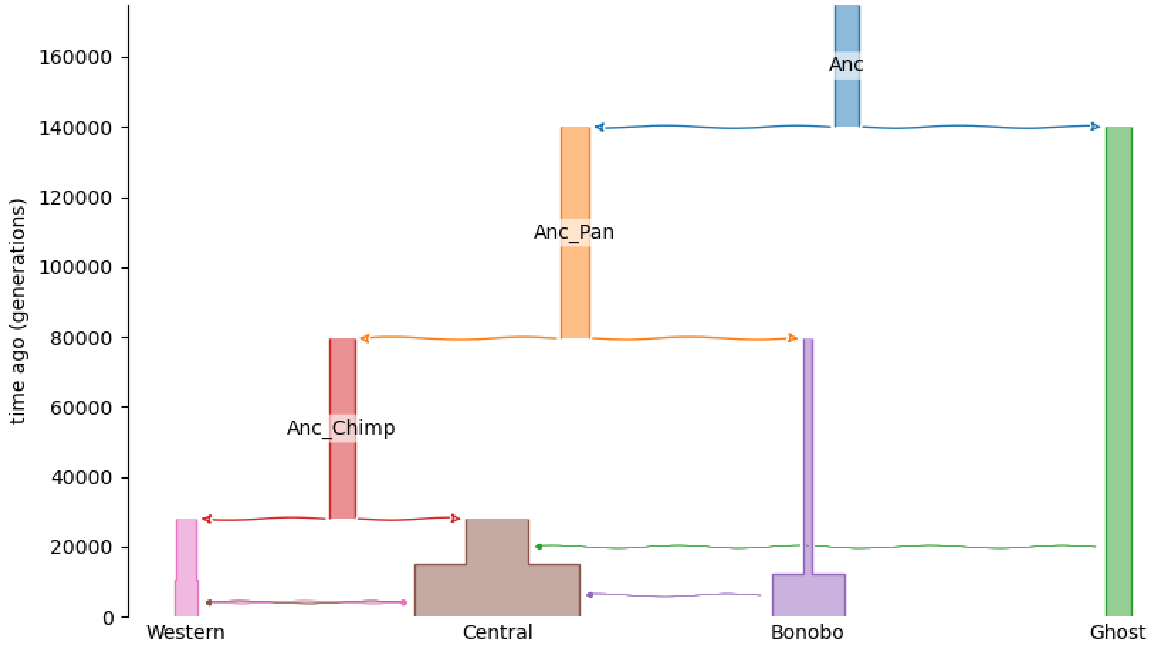


**Figure S5 Chimpanzee-Ghost-Bonobo model for simulating data.** The demographic parameters can be found in Supplementary Table S12.

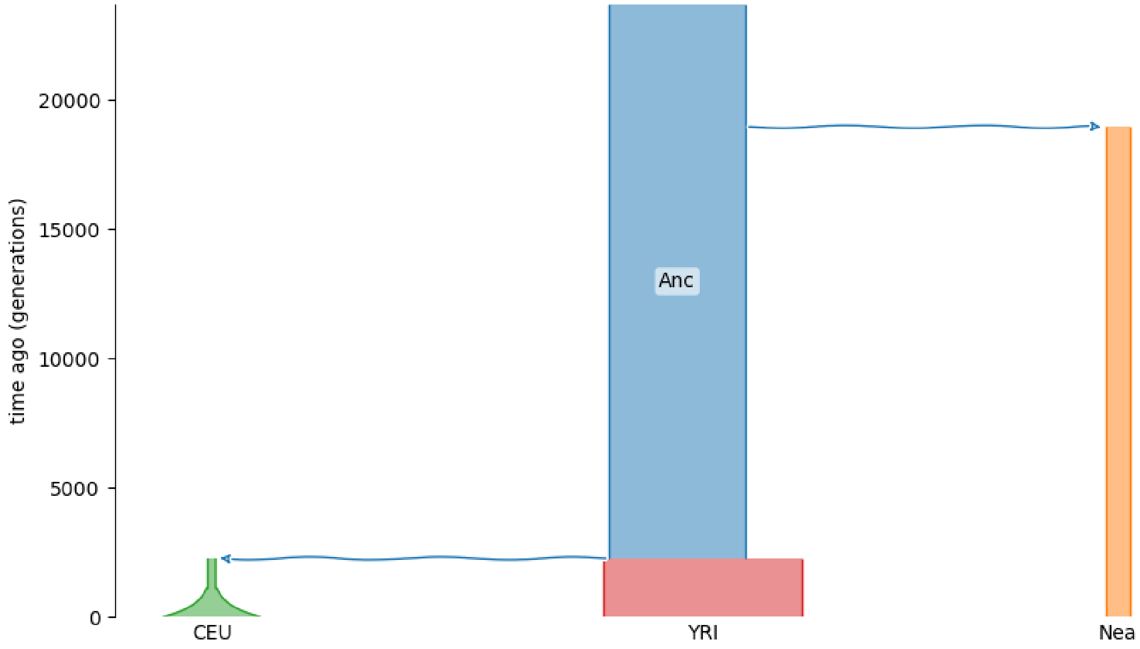


**Figure S6 Human-Neanderthal model without introgression for simulating data to calculate expected *S** scores.** The demographic parameters can be found in Supplementary Table S9.

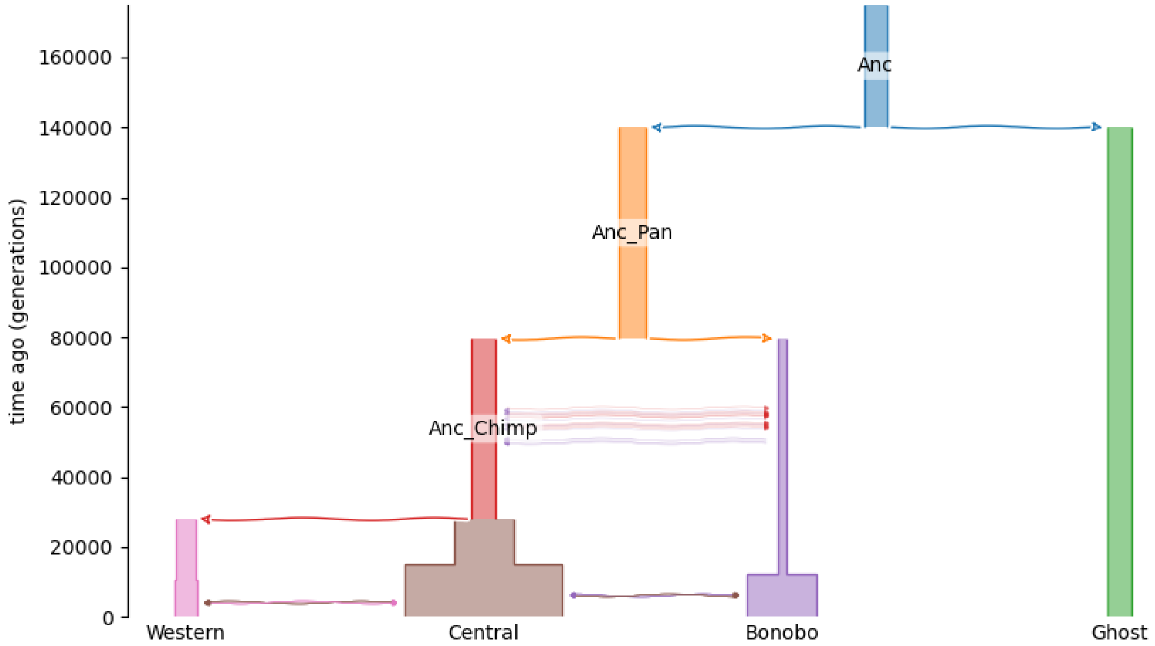


**Figure S7 Bonobo-Ghost model without introgression for simulating data to calculating expected *S** scores.** The demographic parameters can be found in Supplementary Table S10.

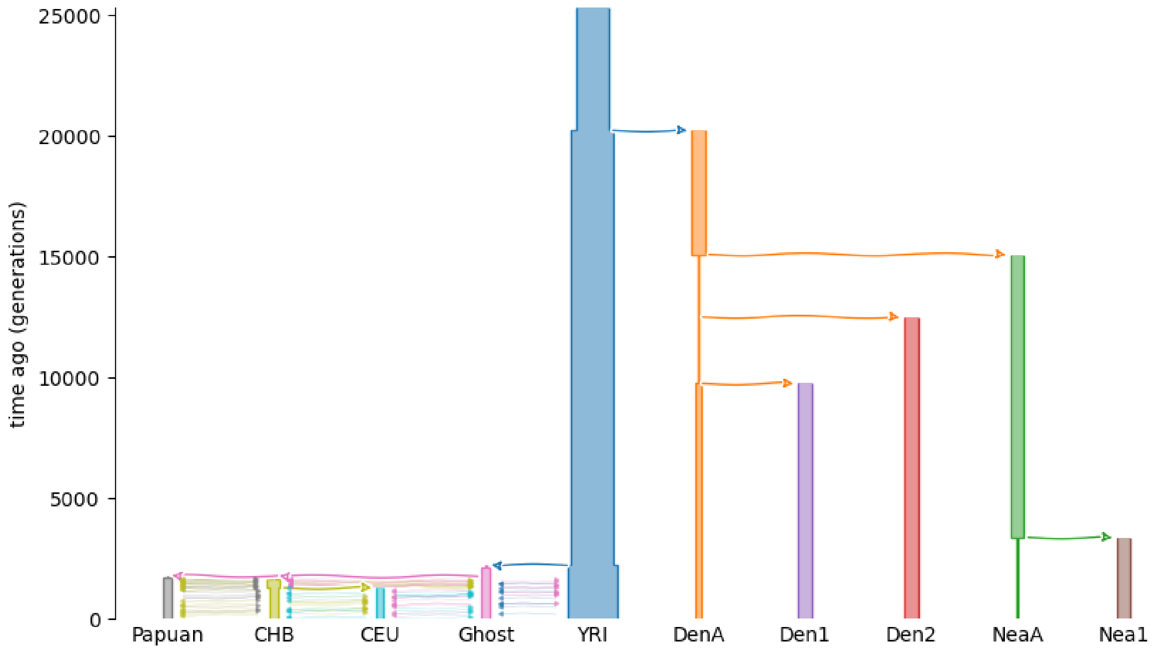


**Figure S8 Human-Neanderthal-Denisovan model without introgression for simulating data to calculate expected *S** scores.** The demographic parameters can be found in Supplementary Table S11.

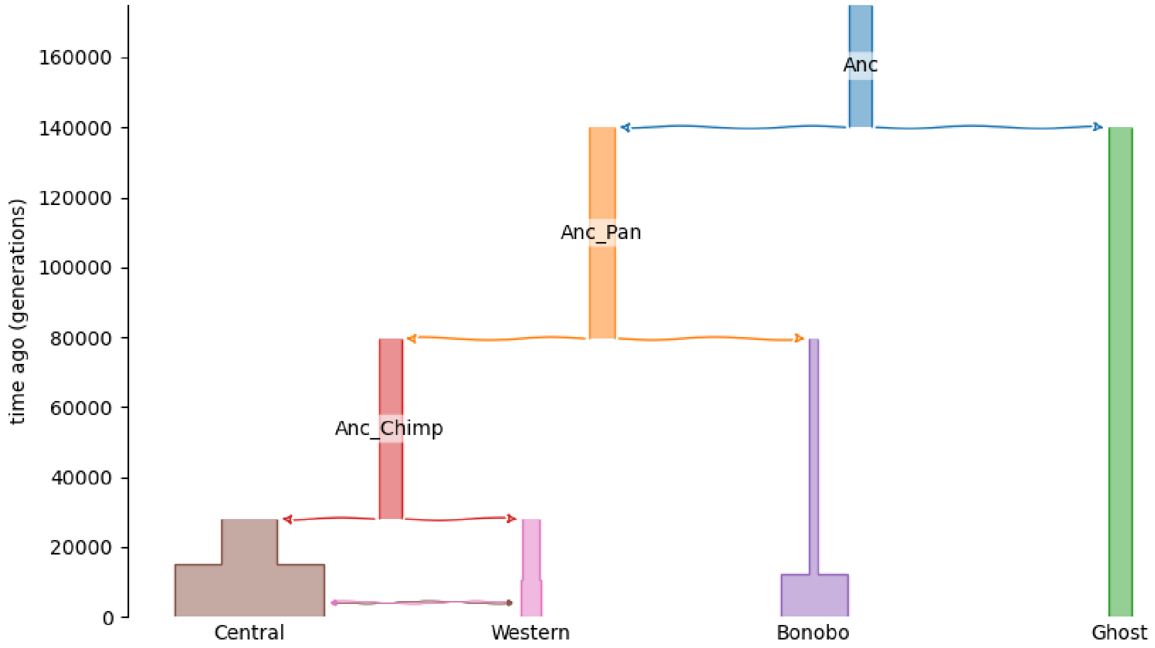


**Figure S9 Chimpanzee-Ghost-Bonobo model without introgression for simulating data to calculate expected *S** scores.** The demographic parameters can be found in Supplementary Table S12.

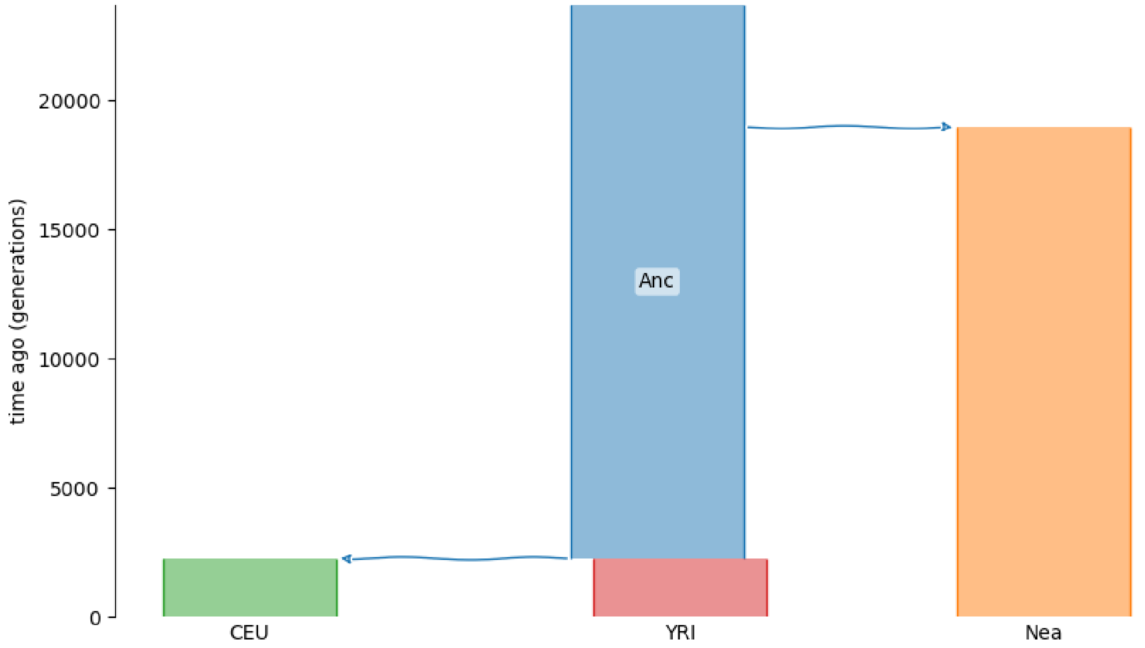


**Figure S10 Constant Human-Neanderthal model without introgression for simulating data to calculate expected *S** scores.** The population size of each population is 10,000. Other demographic parameters can be found in Supplementary Table S9.

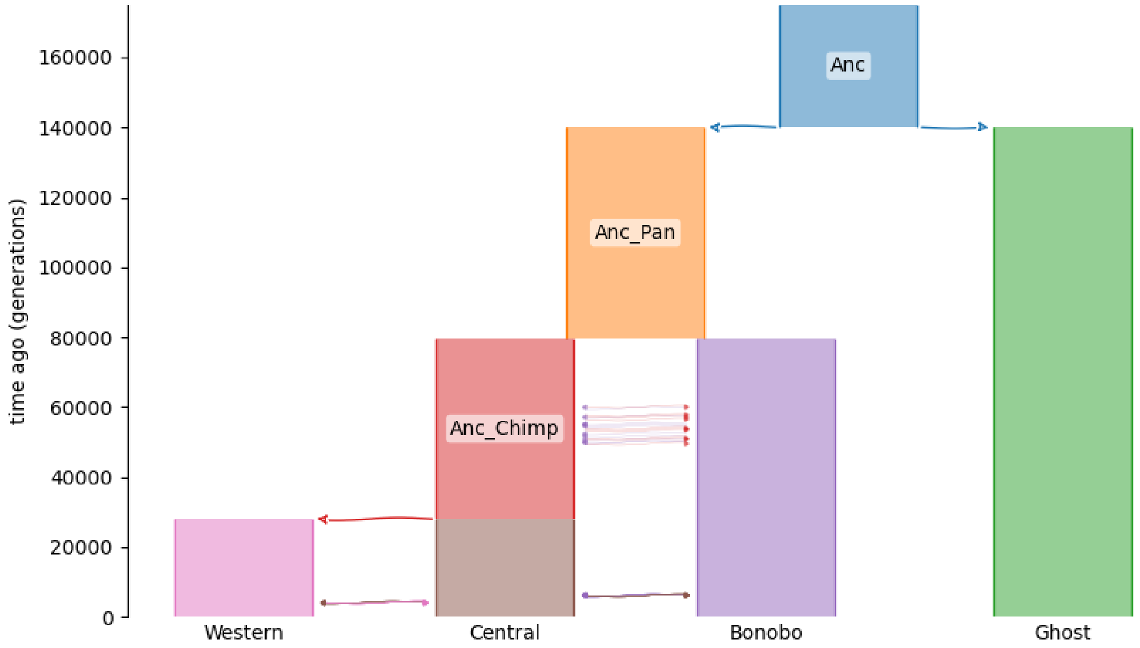


**Figure S11 Constant Bonobo-Ghost model without introgression for simulating data to calculate expected *S** scores.** The population size of each population is 10,000. Other demographic parameters can be found in Supplementary Table S10.

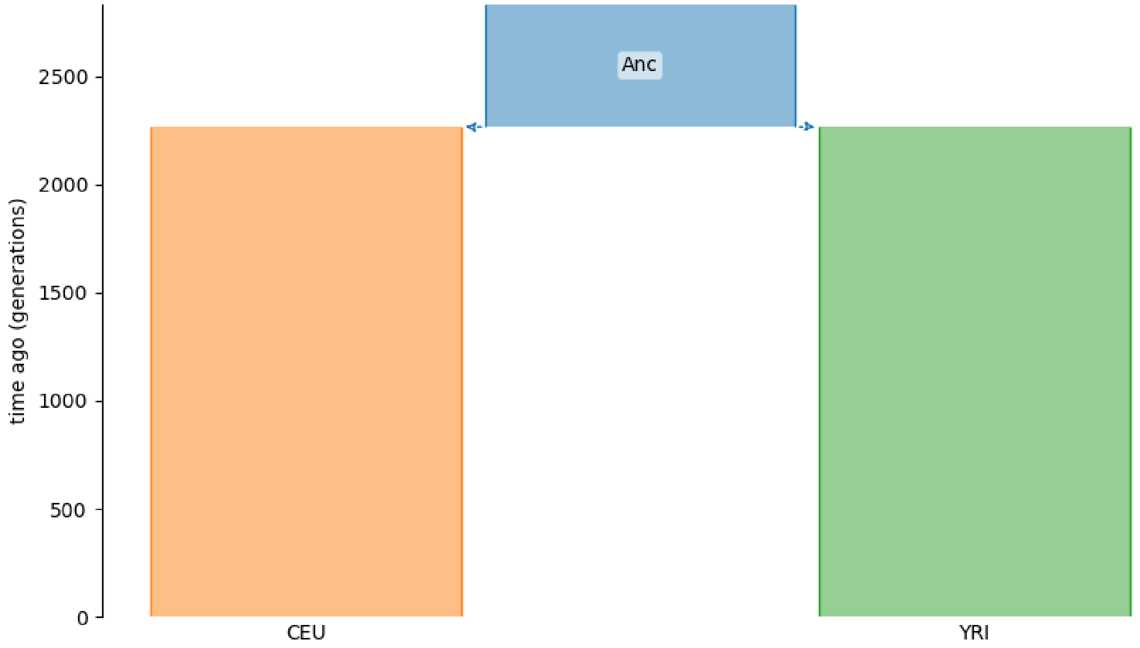


**Figure S12 Constant Human-Neanderthal model using the reference and target populations only for simulating data to calculate expected *S** scores.** The population size of each population is 10,000. The source population (Neanderthal) is not simulated. Other demographic parameters can be found in Supplementary Table S9.

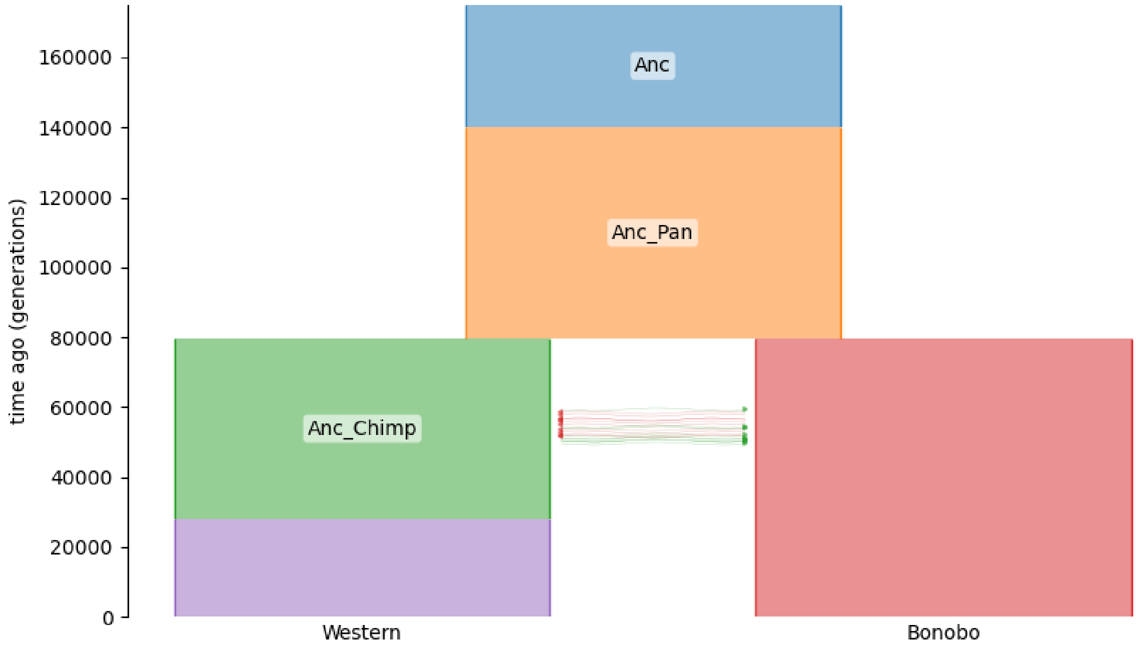


**Figure S13  Constant Bonobo-Ghost model using the reference and target populations only for simulating data to calculate expected *S** scores.** The population size of each population is 10,000. The source population (Ghost) and the Central Chimpanzee population are not simulated. Other demographic parameters can be found in Supplementary Table S10.

### Supplementary Tables

**Table S1 Performance of sstar for detecting ghost introgression (Figure 1B and 1D)**

**Table S2 Performance of SPrime for detecting ghost introgression (Figure 1B and 1D)**

**Table S3 Performance of SkovHMM for detecting ghost introgression (Figure 1B and 1D)**

**Table S4 Performance of sstar for detecting two-source introgression (Figure 1F and 1H)**

**Table S5 Performance of SPrime for detecting two-source introgression (Figure 1F and 1H)**

**Table S6 Performance of ArchaicSeeker2.0 for detecting two-source introgression (Figure 1F and 1H)**

**Table S7 PR AUC of sstar, SPrime and SkovHMM for detecting ghost introgression (Figure 1D and 1E)**

**Table S8 PR AUC of sstar, SPrime and ArchaicSeeker2.0 for detecting two-source introgression (Figure 1G and 1I)**

**Table S9 Parameters for the Human-Neanderthal model (Supplementary Figure S2, S6, S10 and S12)**

**Table S10 Parameters for the Bonobo-Ghost model (Supplementary Figure S3, S7, S11 and S13)**

**Table S11 Parameters for the Human-Neanderthal-Denisovan model (Supplementary S4 and S8)**

**Table S12 Parameters for the Chimpanzee-Ghost-Bonobo model (Supplementary S5 and S9)**
